# LEVERAGING TRANSFER LEARNING FOR HIGH-ACCURACY PHENOTYPIC SCREENING IN ZEBRAFISH IMAGE ANALYSIS

**DOI:** 10.1101/2025.01.21.634041

**Authors:** Varshini Uma Jayaraman, Raghavender Medishetti, Shreetama Ghosh, Kiranam Chatti, Mahesh Kumar Uppada, Srinivas Oruganti

## Abstract

This paper presents a method for classifying zebrafish images captured before and after drug administration. Leveraging the power of transfer learning and fine-tuning, the approach effectively overcomes the challenges of limited datasets in biomedical imaging. By employing a pre-trained convolutional neural network (CNN) as the base model, transfer learning allows us to utilize learned features from large-scale image datasets, significantly reducing training time and computational resources. Fine-tuning specific layers of the model on our zebrafish dataset further enhances its ability to detect subtle visual differences induced by drug administration. The proposed approach achieves high accuracy in classifying zebrafish images, demonstrating its potential as a reliable tool for analysing phenotypic changes due to pharmacological interventions. This model could be instrumental in accelerating drug discovery and research in zebrafish-based assays, offering a scalable and efficient solution for image-based biomedical analysis.

## INTRODUCTION

Zebrafish have emerged as a powerful model organism in toxicity^1,2^ and pharmacological research, primarily due to their unique biological features and suitability for high-throughput screening.^3^ One of the key benefits is their ability to produce large numbers of offspring from a single mating. This high reproductive capacity allows researchers to work with extensive sample sizes, thereby enhancing their experiments statistical power and reliability. Another significant advantage of zebrafish is their transparency during early developmental stages.^4^ This trait provides a remarkable opportunity for in vivo imaging,^5^ allowing researchers to observe internal structures and organ development in real-time. The transparency eliminates the need for invasive procedures, facilitating continuous, non-destructive monitoring of zebrafish throughout various stages of development. This capability is particularly useful for detecting organ-specific toxicities, developmental abnormalities, and phenotypic changes resulting from exposure to experimental compounds.^6^ The rapid development of zebrafish larvae, with the formation of key organ systems within the first 24 to 48 hours post-fertilization, makes them particularly effective for assessing the early effects of compounds on organogenesis and overall development.^7^ Moreover, zebrafish are ideally suited for high-throughput screening, allowing the evaluation of large numbers of compounds in a short period.

Automated imaging systems can capture large datasets from hundreds of embryos or larvae simultaneously, accelerating the process and providing high-volume data for analysis. ^8^ However, despite the promise of high-throughput systems, the available high-throughput screening approaches can be costly, which may limit their accessibility for some research groups. This financial constraint may also affect the ability to implement large-scale screening projects, especially in resource-limited settings. In the context of evaluating the toxicity of a small molecule tyrosine kinase inhibitor,^9^ zebrafish were imaged from the lateral view to assess phenotypic changes such as structural abnormalities, pigmentation alterations, and edema. ^10,11^ Zebrafish imaging is indispensable in the early detection of phenotypic changes, which facilitates the identification of toxicity effects at critical early stages of development. This approach is particularly effective for observing organ-specific toxicities, developmental delays, and monitoring hatching rates, offering a comprehensive view of how a compound affects zebrafish growth and development. ^12^

Traditionally, manual scoring methods have been employed to assess these phenotypic changes, where specific features are evaluated and scored based on predefined criteria. However, manual scoring can be time-consuming, subject to observer bias, and prone to variability, which may lead to inconsistent results. Moreover, subtle or complex phenotypic changes, such as those affecting organ morphology or developmental milestones, can easily be overlooked using manual methods. Although some image analysis tools like ImageJ are widely used for processing zebrafish images, they come with several limitations. These tools, while effective for basic analyses, often struggle with handling large datasets, performing complex image segmentation, or detecting subtle phenotypic changes with high precision. Furthermore, these programs are typically user-dependent, which can introduce inconsistencies in results, especially when different users analyze the same dataset.

Recent advancements in image analysis using artificial intelligence (AI) and machine learning (ML) have significantly enhanced the ability to analyse images in an efficient and unbiased manner. The large volume of image data captured at various developmental stages can now be processed using AI/ML algorithms, facilitating the detection of early phenotypic markers of toxicity that might otherwise go unnoticed.^13,14^ Techniques like convolutional neural networks (CNNs) are particularly useful for extracting intricate features from zebrafish images, and supervised learning models can be trained to recognize specific toxicity-related changes in organs or developmental processes.^15^ Furthermore, AI-powered image segmentation^16^ and pattern recognition algorithms allow for precise quantification of morphological changes, such as organ malformations or delayed hatching, improving both the accuracy and reproducibility of toxicity assessments.^17^

By integrating zebrafish image data sets with AI/ML approaches, researchers can significantly enhance the efficiency and precision of toxicity screening. AI/ML methods offer an advanced toolset for high-throughput screening, enabling the early identification of developmental disruptions and organ-specific toxicities with greater accuracy than traditional manual methods or basic image analysis tools. This capability is particularly valuable for large-scale drug development and safety testing, where it can reduce the time and cost of identifying potential toxicities. Ultimately, leveraging AI/ML technologies to analyze zebrafish phenotypic data not only improves the overall quality and consistency of the results but also enhances the capacity for early detection of toxic effects, making zebrafish a powerful tool in both preclinical research and drug discovery.

## METHODOLOGY

This study employed a structured methodology integrating state-of-the-art techniques in deep learning, with a focus on transfer learning and fine-tuning, to develop classification models for zebrafish images. Below, we detail the steps involved:

### Data Collection

The study began with the acquisition of zebrafish images, including images of zebrafish taken before and after drug administration.

All zebrafish experiments were performed at the zebrafish facility at Dr. Reddy’s Institute of Life Sciences (CPCSEA Registration number 1100/po/Re/s/07/CPCSEA), Hyderabad, India, with approval of the Institutional Animal Ethics Committee. The experiments were performed per the Guidelines for Experimentation on Fishes, 2021, published by CPCSEA.

The zebrafish used were an Indian wild-caught strain bred in-house for at least 2 years. Adult fish were maintained in a standalone housing system (ZebTec, Tecniplast). The system continuously circulates filtered and aerated water to all the fish tanks and regulates temperature (28 ± 1°C), conductivity (450–550 μS), and pH (6.7–7.5). A light-dark cycle of 14 h light and 10 h dark was maintained in the facility. The fish were fed three times daily with a varied diet consisting of dry pellets and live freshly hatched brine shrimp (hatched from frozen artemia cysts, INVE Aquaculture).

Fish over 3 months of age were used for spawning. Sloping breeding tanks of 1.7 L capacity (Tecniplast) with a divider were used for breeding. Pairs of male and female fish (1:2 male to female ratio, typically three males and six females) were kept in the breeding tanks on either side of a divider in the evening before the breeding day. Zebrafish are photoperiodic in their breeding and produce embryos at first light. Breeding pairs typically produce 100–300 embryos in our laboratory, with approximately 80% of them being fertilized. Unfertilized eggs were discarded immediately while fertilized eggs were transferred into a 95-mm Petri dish and washed several times in system water and maintained in a 28°C incubator (Shel Lab) until further use.

Embryos at 4 h postfertilization (hpf) were distributed in 24-well plates in E3 medium and exposed to 10 μM of the anti-cancer drug cabozantinib along with vehicle-treated control (0.1% DMSO in E3 medium). The concentration was selected to represent an LC10 value (1/10th of the expected lethal concentration). Twenty embryos were used per experiment and the study was repeated three times. The treatment was continued for up to 5 days postfertilization (5 dpf) in an incubator at a constant temperature of 28°C (with a photoperiod of 14 h light:10 h dark).

Bright-field microscopic images of the larvae were acquired in a well-lit room under daylight conditions, at an ambient temperature of 28°C, using a Zeiss Stereo Discovery V8 zoom stereo microscope (lit from below by white light) with a ProRes colour camera C3 (3 Megapixels) attached to the C-mount at a resolution of 480X480 pixels.

### Dataset Composition

The dataset was curated from various sources, ensuring that it had a balanced representation of both classes (pre- and post-drug administration). The dataset was split into training, validation, and test sets. Each of the two data sets contained 2 classes namely control (zebrafish images before the administration of the drug) and test (zebrafish images after the administration of the drug).

### Data Preprocessing

The final dataset included images of zebrafish embryos at different stages (4 hpf to 5 dpf) treated with cabozantinib or control medium. The dataset used in this paper comprised two classes: Control and Test, with a total of 451 zebrafish images. For the Control class, 360 images were available, of which 325 were allocated for training, 5 for validation, and 30 for testing. For the Test class, 91 images were available, with 79 used for training, 5 for validation, and 7 for testing. The dataset distribution was carefully chosen to ensure sufficient representation of each class during the training phase, while also setting aside a portion for testing to evaluate the model’s generalization.

Preprocessing was a critical step in preparing the data for deep learning models, particularly to ensure compatibility with pre-trained architectures like ResNet and VGG. The images were resized to a standardized input size of 224×224 pixels, maintaining the aspect ratio to preserve the morphological integrity of the zebrafish specimens. This resizing step ensured uniformity and facilitated effective feature extraction by the models.

To further enhance the consistency of inputs, pixel values were normalized to the [0, 1] range by dividing each value by 255. This normalization improved numerical stability during model training, enabling the network to converge more efficiently. Additionally, dynamic data augmentation techniques were employed during the training phase to increase the variability in the dataset, mitigate overfitting, and enhance the generalization capability of the model.

These augmentation methods included random rotations, horizontal and vertical flips, zooming in and out, and translations along the x and y axes, all of which contributed to a more robust and versatile training process.

### Model selection and transfer learning

Pre-trained deep learning models, including ResNet50, VGG16, and VGG19, were evaluated for their efficacy in zebrafish image classification due to their well-established performance in similar tasks. These models were chosen for their ability to effectively extract features from images and their versatility in handling complex classification problems.

The transfer learning process was implemented to fine-tune these models for zebrafish-specific classification tasks. The initial layers of the pre-trained models, which are designed to capture general image features such as edges and textures, were frozen to retain the knowledge gained from large-scale training on datasets like ImageNet. This prevented the need to re-learn basic visual features and reduced computational requirements. Subsequently, the output layer was replaced with a custom classification head tailored for the specific task. This head consisted of dense layers and a sigmoid activation function to enable binary classification. The newly added layers were then trained using the zebrafish dataset, allowing the model to adapt the pre-learned features to domain-specific patterns and characteristics unique to zebrafish images.

### Fine-Tuning for Optimization

To further enhance model performance, fine-tuning was employed by unfreezing select deeper layers of the pre-trained models. This approach facilitated domain-specific feature learning while preserving the general learned representations from large-scale datasets like ImageNet. By selectively training deeper layers, the models were able to adapt more effectively to the unique characteristics of the zebrafish dataset.

Several optimization techniques were implemented to ensure efficient and effective training. A lower learning rate was utilized during the fine-tuning process to prevent catastrophic forgetting of previously learned features. Dynamic learning adjustments, such as learning rate decay and adaptive optimization strategies like the Adam optimizer, were employed to ensure stable and precise weight updates throughout training. Additionally, various batch sizes and numbers of training epochs were tested to strike a balance between model convergence and computational efficiency. Cross-validation was conducted to identify the optimal combination of hyperparameters, ensuring that the models achieved robust performance across the zebrafish dataset.

### Model Training and Validation

The models were trained on the processed zebrafish dataset, with dynamic data augmentation applied during training to enhance generalization and improve robustness to variations in the data. The binary cross-entropy loss function was minimized during backpropagation to optimize the model’s predictions for the binary classification task. To further prevent overfitting, regularization techniques, including dropout, were incorporated into the training process, ensuring more generalized feature learning.

Model validation was carried out using a separate validation set to monitor performance throughout training. This approach ensured that the model did not overfit to the training data while maintaining a focus on real-world applicability. Key performance metrics, including training and validation accuracy, as well as training and validation loss, were computed after each epoch. These metrics provided insights into the model’s learning progress and guided adjustments to improve its performance and reliability.

### Model Evaluation and Testing

The best-performing model, selected based on its superior validation performance, was rigorously evaluated on an independent test set to assess its accuracy on unseen data. Accuracy, defined as the proportion of correctly classified images, was used as the primary evaluation metric to quantify the model’s effectiveness in real-world scenarios.

### Comparative Analysis of Models

A comparative analysis was conducted among the pre-trained models, including ResNet50, VGG16, and VGG19. These models were evaluated based on classification accuracy, computational efficiency, and generalizability to determine their suitability for zebrafish image classification. The final model was selected for its ability to achieve a balance between high classification performance and resource efficiency, ensuring its practicality for deployment in research or diagnostic settings.

## RESULTS AND DISCUSSION

### Model Performance Overview

The classification models, developed using transfer learning and fine-tuning techniques, demonstrated strong efficacy in distinguishing between zebrafish images captured before and after drug administration. Despite the modest dataset size, the use of transfer learning and fine-tuning mitigated the challenges typically associated with limited data availability. Transfer learning enabled the model to leverage pre-trained weights from large-scale datasets, extracting meaningful features that generalized well to zebrafish phenotypic analysis. This approach ensured robust classification despite the dataset’s size constraints.

Pre-trained models including ResNet50, VGG16 and VGG19, were evaluated for their performance, highlighting the benefits of transfer learning combined with fine-tuning in enhancing model capabilities.

### Quantitative Results

Among the models tested, VGG19 emerged as the best-performing model after fine-tuning. The key metrics for VGG19 were as follows:

Accuracy: 0.9759

Loss: 0.1083

Validation Accuracy: 0.9000

Validation Loss: 0.1633

### Comparison of Models

A comparative analysis was conducted to evaluate the performance of a simple self-built classification model alongside pre-trained models, including ResNet-50, VGG-16, and VGG-19, after fine-tuning. The results, summarized in **Table 1**, highlight the classification accuracy, loss, validation accuracy, and validation loss achieved by each model. This analysis underscores the differences in model performance and generalization capabilities for the zebrafish classification task.

**Table 1.** Performance metrics of models.

| MODELS | ACCURACY | LOSS | VALIDATION<br>ACCURACY | VALIDATION<br>LOSS |
| --- | --- | --- | --- | --- |
| Simple<br>Classifier | 0.9729 | 10.1984 | 0.8 | 1.3157 |
| RESNET-50 | 0.7625 | 0.5506 | 0.5 | 0.8253 |
| VGG16 | 0.8578 | 0.339 | 0.8 | 0.3689 |
| VGG19 | 0.9759 | 0.1083 | 0.9 | 0.1633 |

The Simple Classifier achieved strong training accuracy (0.9729) and moderate validation accuracy (0.8), but its high training loss (10.1984) and validation loss (1.3157) indicate potential overfitting or suboptimal convergence. ResNet-50, despite its deep architecture, struggled with both training and validation metrics, achieving only 0.7625 accuracy and 0.5 validation accuracy, along with relatively high loss values. This suggests limited adaptability to the zebrafish dataset and a need for further fine-tuning. VGG-16 demonstrated improved performance compared to ResNet-50, achieving a training accuracy of 0.8578 and validation accuracy of 0.8. Its training and validation losses (0.339 and 0.3689, respectively) were significantly lower, reflecting better generalization. However, fluctuations in its validation trends suggest some instability in learning.

VGG-19 emerged as the best-performing model, achieving the highest training (0.9759) and validation accuracy (0.9). Its training and validation losses (0.1083 and 0.1633) were the lowest among all models, indicating effective learning and strong generalization. The steady improvement in both accuracy and loss during training and validation, as depicted in **Figure 4**, further emphasizes its suitability for the zebrafish classification task.

**Figures 1** through **4** illustrate the training and validation accuracy and loss curves for the evaluated models. The Simple Classifier (**Figure 1**) showed a steady increase in training accuracy but lacked validation metrics beyond the fifth epoch. ResNet-50 (**Figure 2**) exhibited fluctuating training accuracy and high validation loss, signalling insufficient generalization. VGG-16 (**Figure 3**) displayed more consistent improvements but with some variability in validation loss. VGG-19 (**Figure 4**) stood out with stable, converging trends in both accuracy and loss, making it the most robust and reliable model for this study.

**Figure 1.**
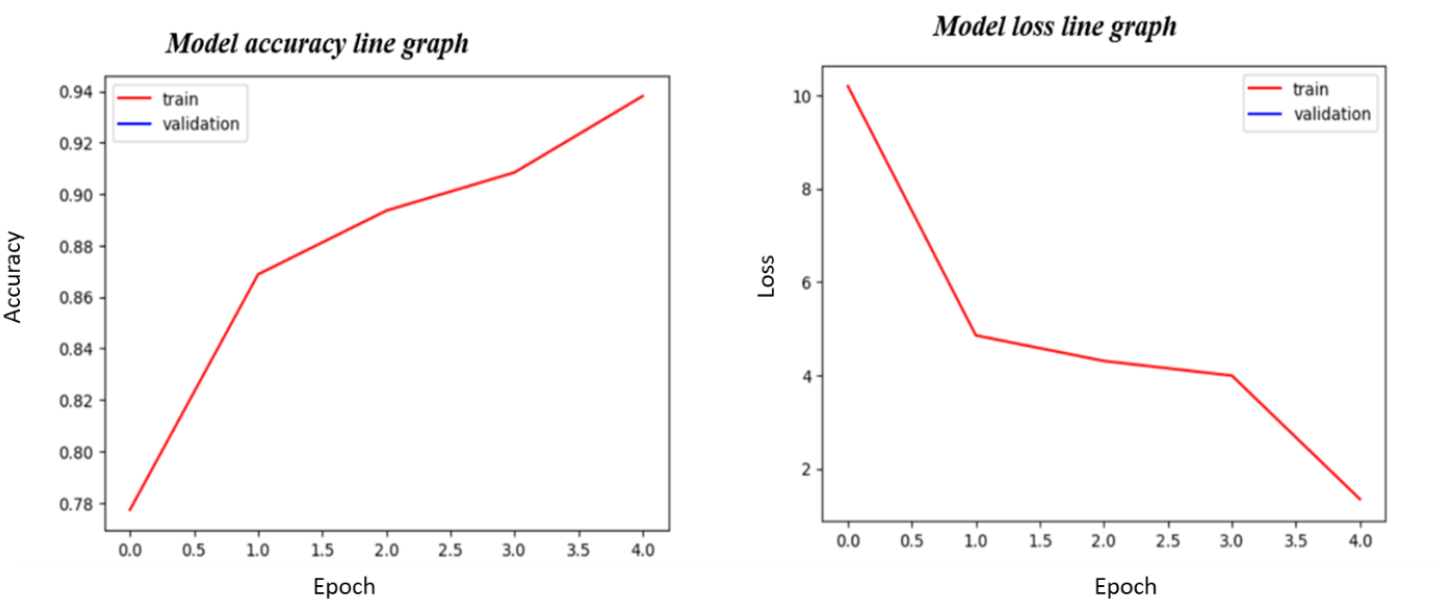
Model results for simple classifier model after training the model at the most of 5 epochs with a dataset of 404 augmented images in 2 classes

**Figure 2.**
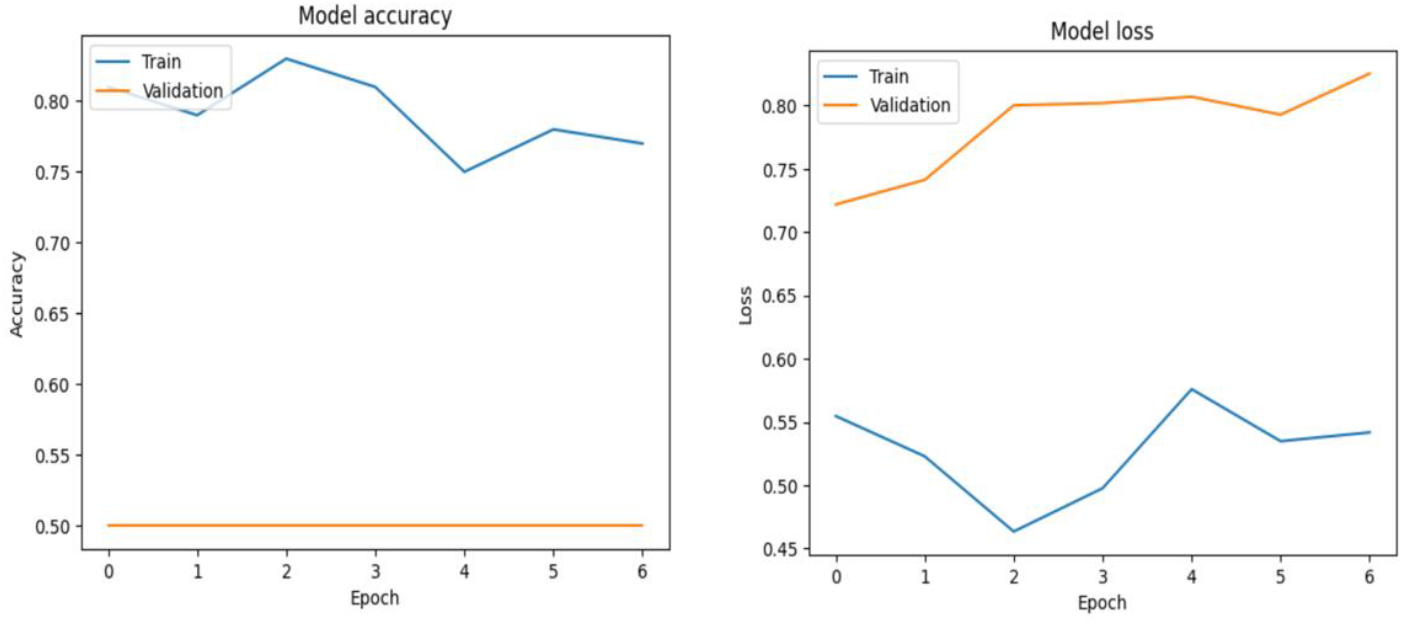
Model results for RESNET-50 after training and fine tuning the model at the most of 7 epochs with a dataset of 404 augmented images in 2 classes

**Figure 3.**
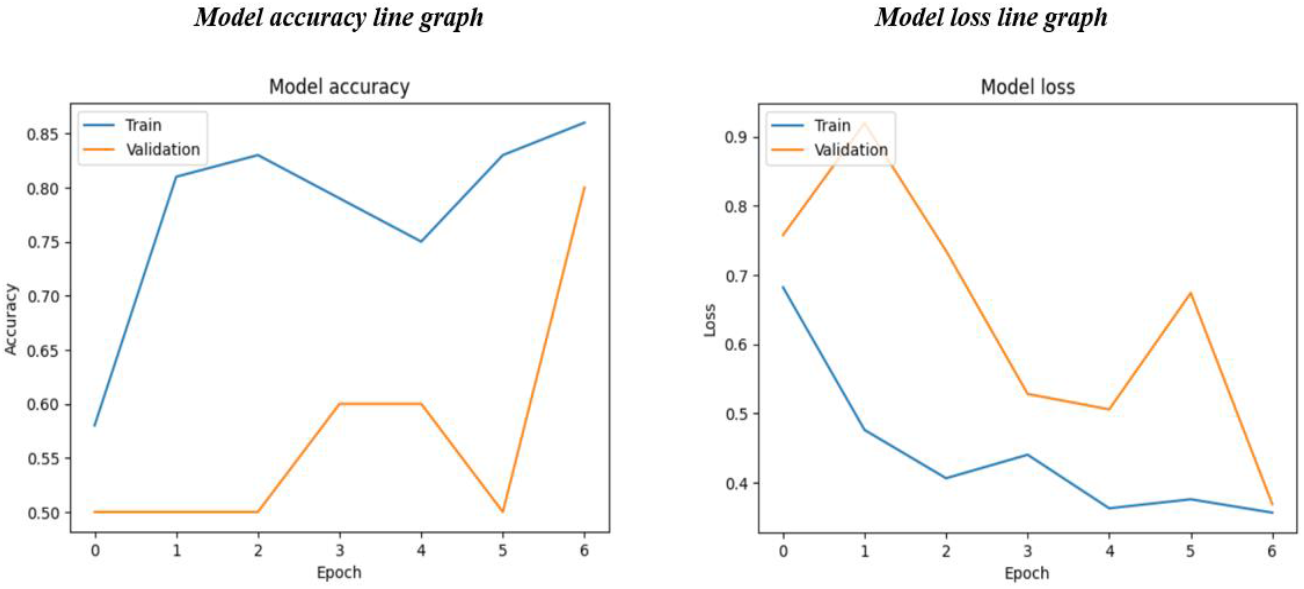
Model results for VGG-16 after training and fine-tuning the model at the most of 7 epochs with a dataset of 404 augmented images in 2 classes

**Figure 4.**
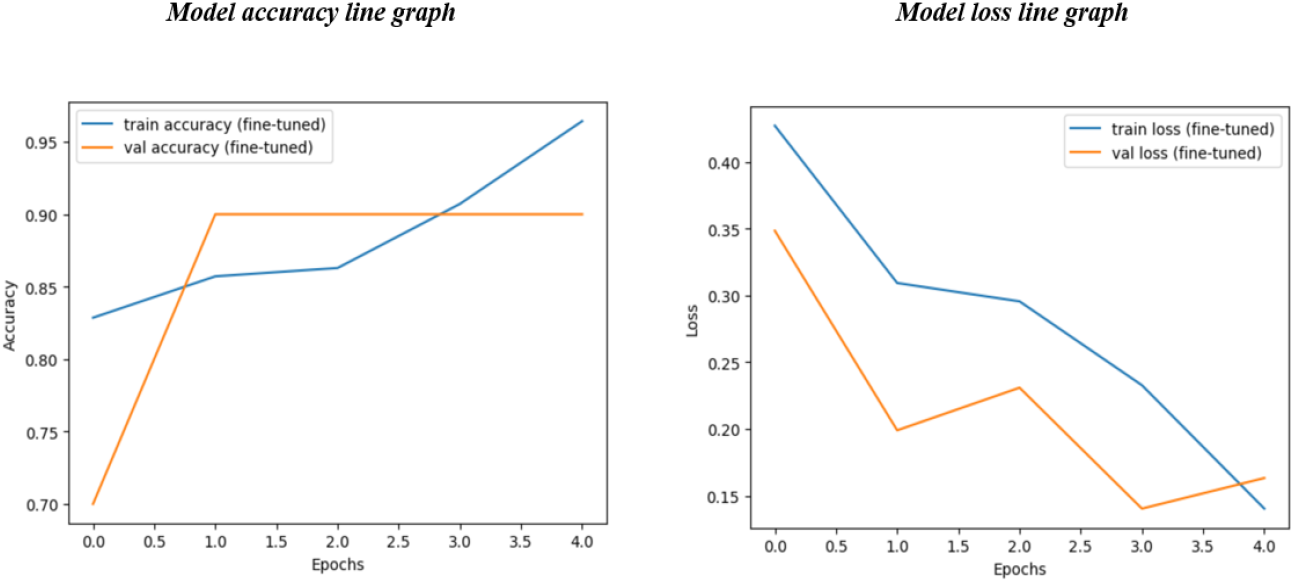
Model results for VGG-19 after training and fine tuning the model at the most of 5 epochs with a dataset of 404 augmented images in 2 classes

In conclusion, the comparative analysis highlights VGG-19 as the optimal choice for zebrafish classification, outperforming other models in accuracy, loss reduction, and overall generalization capability.

### Model Inference

The inference results for all models, visualized in **Figures 5–8**, demonstrate their ability to classify zebrafish images into control (Class 0) and drug-administered (Class 1) categories. The Simple Classifier (**Figure 5**) struggled with generalization, misclassifying both test samples and assigning incorrect labels. ResNet-50 (**Figure 6**) showed slight improvement, correctly classifying one control image but mislabelling the drug-administered sample, reflecting its limited adaptability. VGG-16 (**Figure 7**) demonstrated better performance, correctly identifying both control and drug-administered images in most cases, though occasional errors occurred in more challenging scenarios. VGG-19 (**Figure 8**) stood out as the most robust model, accurately predicting both control and drug-administered images without misclassification, aligning with its high validation performance and low loss observed during training. These results underscore VGG-19’s superior reliability and effectiveness in zebrafish classification tasks.

**Figure 5.**
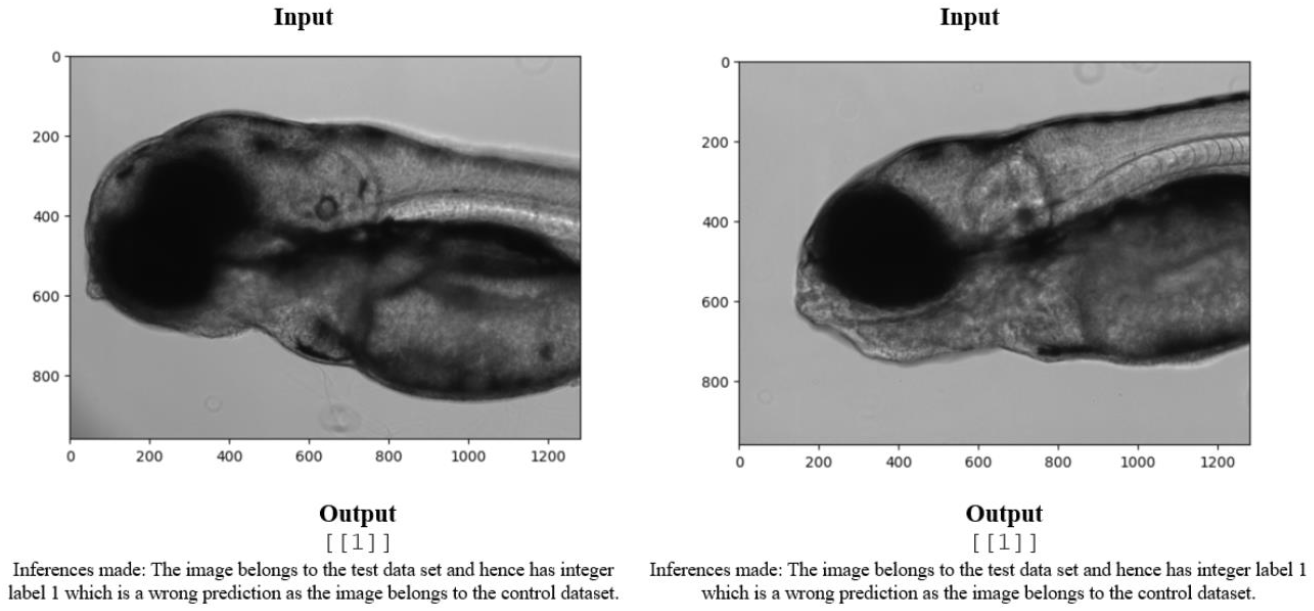
Model inference using simple classifier

**Figure 6.**
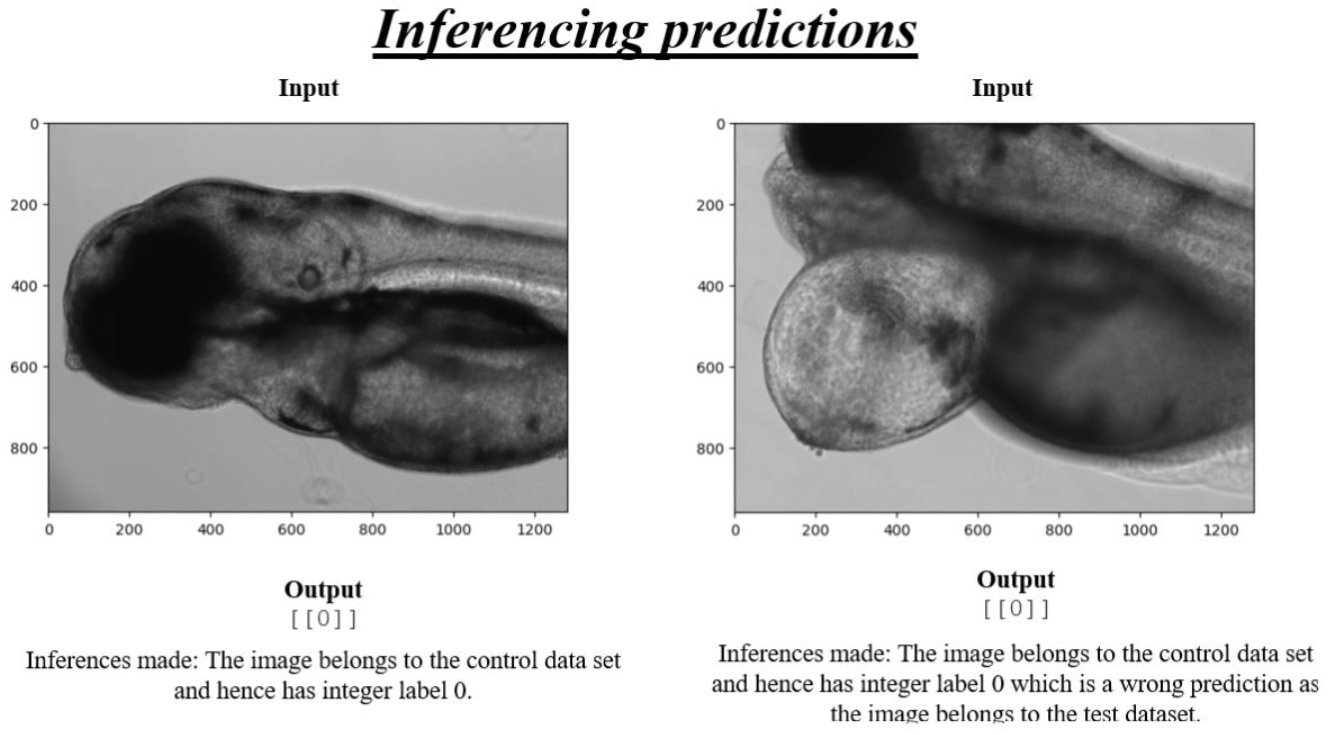
Model inference using RESNET-50

**Figure 7.**
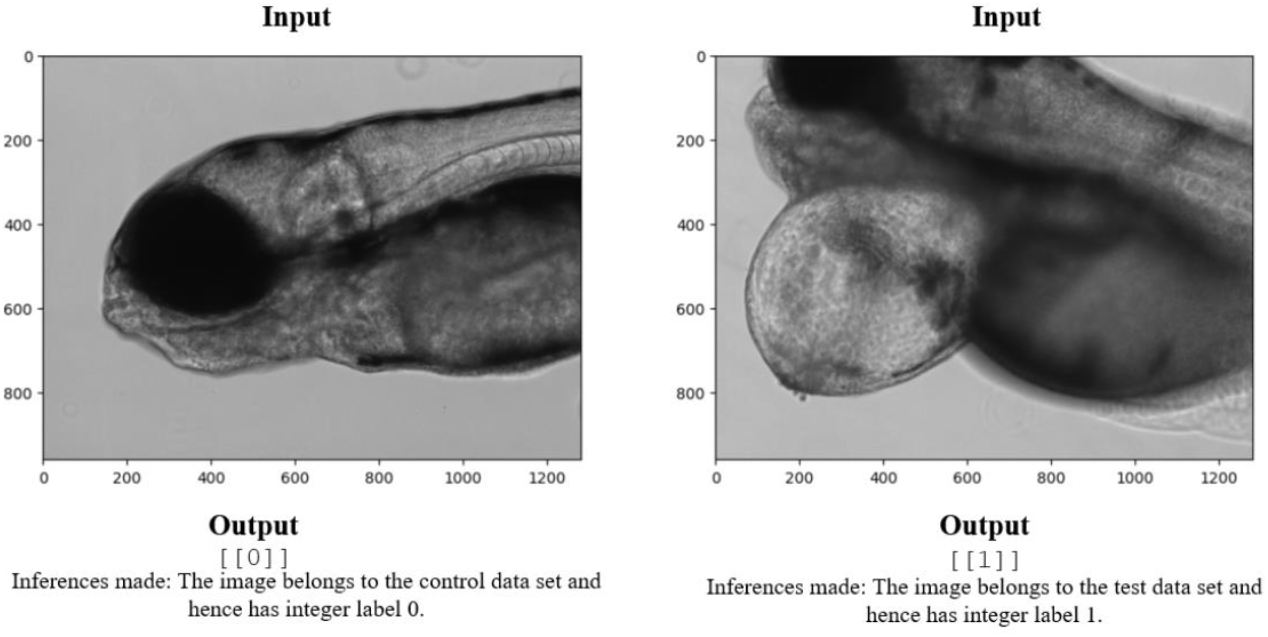
Model inference using VGG16

**Figure 8.**
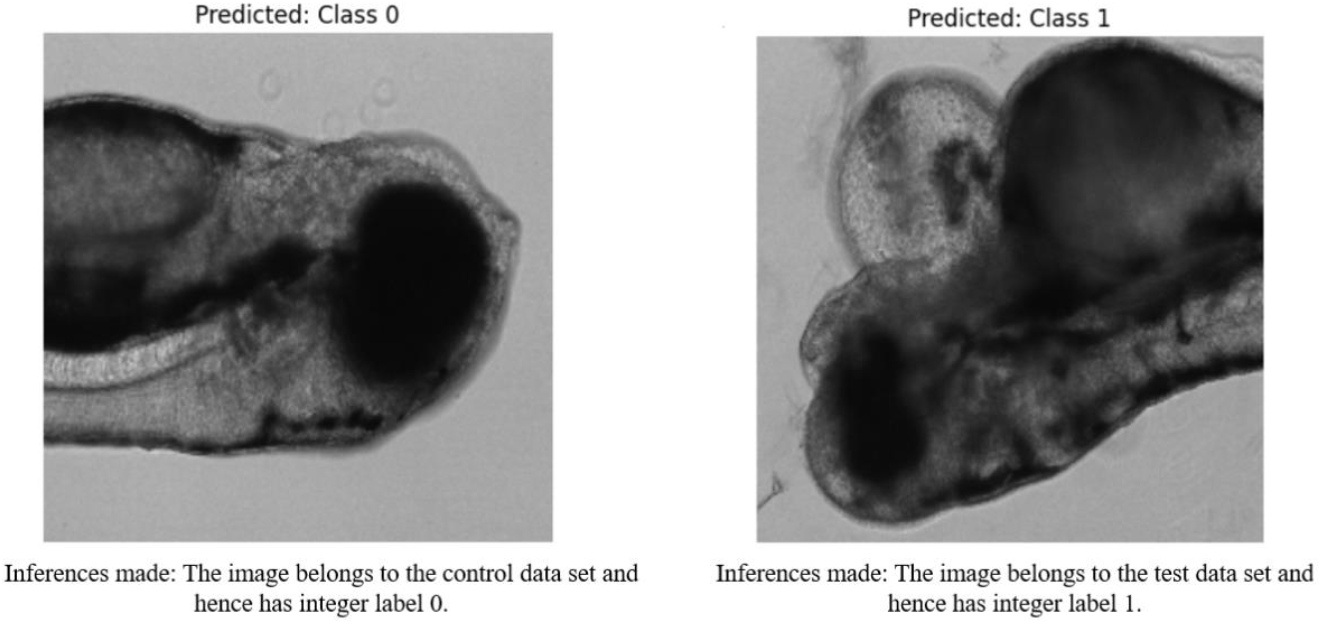
Model inference using VGG19

## CONCLUSION

The study successfully demonstrated that deep learning models using transfer learning and fine-tuning can accurately classify zebrafish images before and after drug administration. The best-performing model, VGG19, achieved an accuracy of 0.9759, demonstrating strong performance in distinguishing subtle differences in zebrafish morphology. Fine-tuning the pre-trained models significantly improved the classification accuracy, boosting performance from an initial 0.7887 (transfer learning only) to over 0.9759. This process allowed the model to focus on zebrafish-specific features induced by drug administration, capturing nuanced changes that were otherwise difficult to detect. One of the main challenges encountered was the imbalance between the number of images of zebrafish before and after drug administration. While data augmentation techniques helped mitigate this issue, an even larger and more balanced dataset could further improve the model’s performance and robustness. The dataset size was very small compared to larger image classification tasks. A larger, more diverse dataset would likely improve model generalization and provide more reliable results across different drugs and zebrafish species. The study highlights the effectiveness of using deep learning, specifically transfer learning and fine-tuning, for classifying zebrafish images before and after drug administration. The high classification accuracy, coupled with the ability to identify drug-induced morphological changes, underscores the potential of this approach for automated drug testing and biomedical research. While challenges such as data imbalance and computational constraints were encountered, the findings suggest that with further improvements in dataset size, segmentation, and model architecture, the models can be refined and extended for broader applications in drug discovery and toxicology. The study showed the potential of using deep learning models in automated drug testing workflows, especially in biomedical research. The model can be extended to identify the effects of various drugs on zebrafish or even other species, making it a valuable tool for large-scale drug testing and toxicology studies.

## ASSOCIATED CONTENT

There is no supporting information for this paper.

## AUTHOR CONTRIBUTIONS

This manuscript was written through contributions of all authors. All authors participated in the development of the study and have given approval to the final version of the manuscript.

## COMPETING INTERESTS

The authors declare no competing financial interest.

## ACKNOWLEDGEMENTS

The authors thank R.I.C.H. for their support.

## ABBREVIATIONS

CNN: Convolutional Neural Network
AI: Artificial Intelligence
VGG: Visual Geometry Group

## Notes

### Competing Interest Statement

The authors have declared no competing interest.

## REFERENCES

1. Cassar S, Adatto I, Freeman JL, Gamse JT, Iturria I, Lawrence C, Muriana A, Peterson RT, Van Cruchten S, Zon LI. Use of Zebrafish in Drug Discovery Toxicology. Chem Res Toxicol. 2020 Jan 21;33(1):95–118. DOI: 10.1021/acs.chemrestox.9b00335. Epub 2019 Nov 16. PMID: 31625720; PMCID: PMC7162671.

2. Sukardi H, Chng HT, Chan EC, Gong Z, Lam SH. Zebrafish for drug toxicity screening: bridging the in vitro cell-based models and in vivo mammalian models. Expert Opin Drug Metab Toxicol. 2011 May;7(5):579–89. DOI: 10.1517/17425255.2011.562197. Epub 2011 Feb 23. PMID: 21345150.

3. Kithcart A, MacRae CA. Using Zebrafish for High-Throughput Screening of Novel Cardiovascular Drugs. JACC Basic Transl Sci. 2017 Feb 27;2(1):1–12. DOI: 10.1016/j.jacbts.2017.01.004. PMID: 30167552; PMCID: PMC6113531.

4. Kimmel CB, Ballard WW, Kimmel SR, Ullmann B, Schilling TF. Stages of embryonic development of the zebrafish. Dev Dyn. 1995 Jul;203(3):253–310. DOI: 10.1002/aja.1002030302. PMID: 8589427.

5. Keller PJ. In vivo imaging of zebrafish embryogenesis. Methods. 2013 Aug 15;62(3):268–78. DOI: 10.1016/j.ymeth.2013.03.015. Epub 2013 Mar 21. PMID: 23523701; PMCID: PMC3907156.

6. Lele Z, Krone PH. The zebrafish as a model system in developmental, toxicological and transgenic research. Biotechnol Adv. 1996;14(1):57–72. DOI: 10.1016/0734-9750(96)00004-3. PMID: 14536924.

7. Zon LI, Peterson RT. In vivo drug discovery in the zebrafish. Nat Rev Drug Discov. 2005 Jan;4(1):35–44. DOI: 10.1038/nrd1606. PMID: 15688071.

8. Spomer W, Pfriem A, Alshut R, Just S, Pylatiuk C. High-throughput screening of zebrafish embryos using automated heart detection and imaging. J Lab Autom. 2012 Dec;17(6):435–42. DOI: 10.1177/2211068212464223. Epub 2012 Oct 10. PMID: 23053930.

9. Medishetti R, C MR, Chatti K. Cabozantinib-induced edema in zebrafish represents an adverse effect characterized by defects in lymphatic vasculature and renal function. J Biochem Mol Toxicol. 2023 Sep;37(9):e23413. DOI: 10.1002/jbt.23413. Epub 2023 Jun 19. PMID: 37335823.

10. Bozhko DV, Myrov VO, Kolchanova SM, Polovian AI, Galumov GK, Demin KA, Zabegalov KN, Strekalova T, de Abreu MS, Petersen EV, Kalueff AV. Artificial intelligence-driven phenotyping of zebrafish psychoactive drug responses. Prog Neuropsychopharmacol Biol Psychiatry. 2022 Jan 10;112:110405. DOI: 10.1016/j.pnpbp.2021.110405. Epub 2021 Jul 25. PMID: 34320403.

11. Planchart A, Mattingly CJ, Allen D, Ceger P, Casey W, Hinton D, Kanungo J, Kullman SW, Tal T, Bondesson M, Burgess SM, Sullivan C, Kim C, Behl M, Padzilla S, Reif DM, Tanguay RL, Hamm J. Advancing toxicology research using in vivo high throughput toxicology with small fish models. ALTEX. 2016;33(4):435–452. DOI: 10.14573/altex.1601281. Epub 2016 Jun 21. PMID: 27328013; PMCID: PMC5270630.

12. Spitsbergen JM, Kent ML. The state of the art of the zebrafish model for toxicology and toxicologic pathology research--advantages and current limitations. Toxicol Pathol. 2003 Jan-Feb;31 Suppl(Suppl):62–87. DOI: 10.1080/01926230390174959. PMID: 12597434; PMCID: PMC1909756.

13. Wang R, Wang B, Chen A. Application of machine learning in the study of development, behavior, nerve, and genotoxicity of zebrafish. Environ Pollut. 2024 Oct 1;358:124473. DOI: 10.1016/j.envpol.2024.124473. Epub 2024 Jun 28. PMID: 38945191.

14. Green AJ, Mohlenkamp MJ, Das J, Chaudhari M, Truong L, Tanguay RL, Reif DM. Leveraging high-throughput screening data, deep neural networks, and conditional generative adversarial networks to advance predictive toxicology. PLoS Comput Biol. 2021 Jul 2;17(7):e1009135. DOI: 10.1371/journal.pcbi.1009135. PMID: 34214078; PMCID: PMC8301607.

15. Jeanray N, Marée R, Pruvot B, Stern O, Geurts P, Wehenkel L, Muller M. Phenotype classification of zebrafish embryos by supervised learning. PLoS One. 2015 Jan 9;10(1):e0116989. DOI: 10.1371/journal.pone.0116989. PMID: 25574849; PMCID: PMC4289190.

16. Ronneberger, O., Fischer, P., & Brox, T. (2015). “U-Net: Convolutional networks for biomedical image segmentation.” International Conference on Medical Image Computing and Computer-Assisted Intervention. DOI: 10.48550/arXiv.1505.04597.

17. Dong G, Wang N, Xu T, Liang J, Qiao R, Yin D, Lin S. Deep Learning-Enabled Morphometric Analysis for Toxicity Screening Using Zebrafish Larvae. Environ Sci Technol. 2023 Nov 21;57(46):18127–18138. DOI: 10.1021/acs.est.3c00593. Epub 2023 Mar 27. PMID: 36971266.

